# Optical Probing of Local Membrane Potential with Fluorescent Polystyrene Beads

**DOI:** 10.1101/2021.04.21.440831

**Authors:** Zehavit Shapira, Nurit Degani-Katzav, Shimon Yudovich, Asaf Grupi, Shimon Weiss

## Abstract

Studying the electrical activity in single cells and in local circuits of excitable cells, like neurons, requires an easy to use and high throughput methodology that enables the measurement of membrane potential. Studying the electrical properties in particular sub-compartments of neurons, or in a specific type of neurons produces additional complexity. An optical voltage-imaging technique that allows high spatial and temporal resolution could be an ideal solution. However, most of the valid voltage imaging techniques are nonspecific; The ones that are more site-directed require much pre-work and specific adaptations in addition to other disadvantages. Here, a new technique for membrane voltage imaging, based on FRET between fluorescent polystyrene (FPS) beads and Dipicrylamine (DPA) is explored. Not only fluorescent intensity is demonstrated to be correlated with membrane potential, but more importantly, single particle voltage detection is demonstrated. Among other advantages, FPS beads can be synthesized with functional surface groups, and be further targeted to specific proteins via conjugation of recognition molecules. Therefore, FPS beads, in the presence of DPA, constitute single-particle detectors for membrane voltage, with a potential to be localized to specific membrane compartments. This new and accessible platform for targeted optical voltage imaging may further elucidate the mechanisms of neuronal electrical activity.

## Introduction

Investigating the electrical properties of excitable cells, such as neurons and muscle cells, is a crucial study in neuroscience. The traditional method in electrophysiology for measurement of membrane potential is the patch-clamp technique. This method provides high resolution electrophysiological recordings from single cells down to single channels. It allows measurements of membrane voltage changes in primary cultured neurons as well as in cultured cell lines. However, a major disadvantage of this method is the use of invasive electrodes which limits the measurements to single cells and results in a low throughput screening assay. A different high throughput assay is needed particularly for electrical measurements from assembly of cells and from different parts of the single neuron itself, such as dendrites and spines. Optical techniques which correlate membrane potential to fluorescent response may deal with these drawbacks; they are relatively non-invasive, providing spatial resolution in assemblies of cells or in sub-cellular regions and are high-throughput.

There are different optical imaging techniques used for the investigation of neuronal activities. A common technology is the calcium imaging [1] method which enables the assessment of the network activity of neurons, based on the intracellular calcium concentration as an indicator of neural activity. Major disadvantages of this method is the lack of information about hyperpolarizing and subthreshold depolarizing signals, relatively low temporal resolution and high background noise.

Voltage-sensitive dyes that enable direct fluorescent imaging of membrane voltage changes are used for neuroimaging at the ensemble level and in both in-vitro and in-vivo formats. However, their potential for neuroimaging is not fully exploited due to shortcomings such as alteration of membrane capacitance, phototoxicity, photobleaching, and nonspecific background labeling. More importantly, they only display non-specific staining. They do not allow for localized, site specific membrane potential measurements.

An interesting technology is the development of genetically encoded voltage indicators (GEVIs) which are engineered fluorescent protein sensors. Indeed, they allow measurements of voltage signals from small neurons and subcellular regions [2], yet detection of action potentials is limited because many suffer from relatively low sensitivity, [3], poor expression [4]and relatively slow kinetics [3, 5, 6].

Recently, voltage sensing nanoparticles (vsNPs) such as voltage sensing quantum dots ([7, 8], have been developed for non-invasive optical recording of membrane potential at the single particle and nanoscale level, at multiple sites, in a large field-of-view. In contrast to voltage-sensitive dyes and GEVIs, vsNPs allow for ‘pointy’, single particle voltage detection. However, vsNPs still face challenges associated with functionalization, bioconjugation, stable membrane insertion, toxicity, and uniformity ([9–13], and additional improvements are needed before this new generation of sensors can be widely translated to neurophysiological applications.

Another interesting mechanism of optical voltage sensors is based on Förster resonance energy transfer (FRET) between fixed fluorescent donors to translocating oxonol acceptors [14]. DiO (Dioctadecyloxacarbocyanine)/DPA (Dipicrylamine) is a known FRET-based membrane potential detection system. DiO is a nontoxic fluorescent lipophilic dye that readily labels membranes. It serves as a stationery FRET donor in the plasma membrane. DPA serves as a mobile FRET acceptor. DPA is a non-fluorescent lipophilic anion which partitions between leaflets of the lipid bilayer in response to membrane potential, thus acting as a voltage sensor.

Our theme was to use this mechanism in order to develop a single particle voltage-detecting system which will enable membrane potential measurements in localized areas. To achieve this goal, nanoparticles were considered to be used as the stationery FRET donors instead of DiO. Fluorescent polystyrene (FPS) beads were identified as the potential candidates. These beads are commercially available in a range of uniform sizes (down to ~20 nm in diameter) and can be loaded with dyes spectrally suitable for FRET interaction with DPA.

The potential for FPS beads to act as voltage sensing nanoparticles, is also attributed to the fact that FPS beads typically have a very thin layer of surface groups which brings the surface of the FPS beads to practically direct contact with the plasma membrane. Since the fluorophores encapsulated within the FPS bead are homogenously distributed, the population of fluorophores closest to the membrane surface may FRET with DPA distributed in the plasma membrane. Therefore, FPS beads with an emission spectral that overlaps with the absorbance of DPA were selected (Fig.1A).

**Figure 1:**
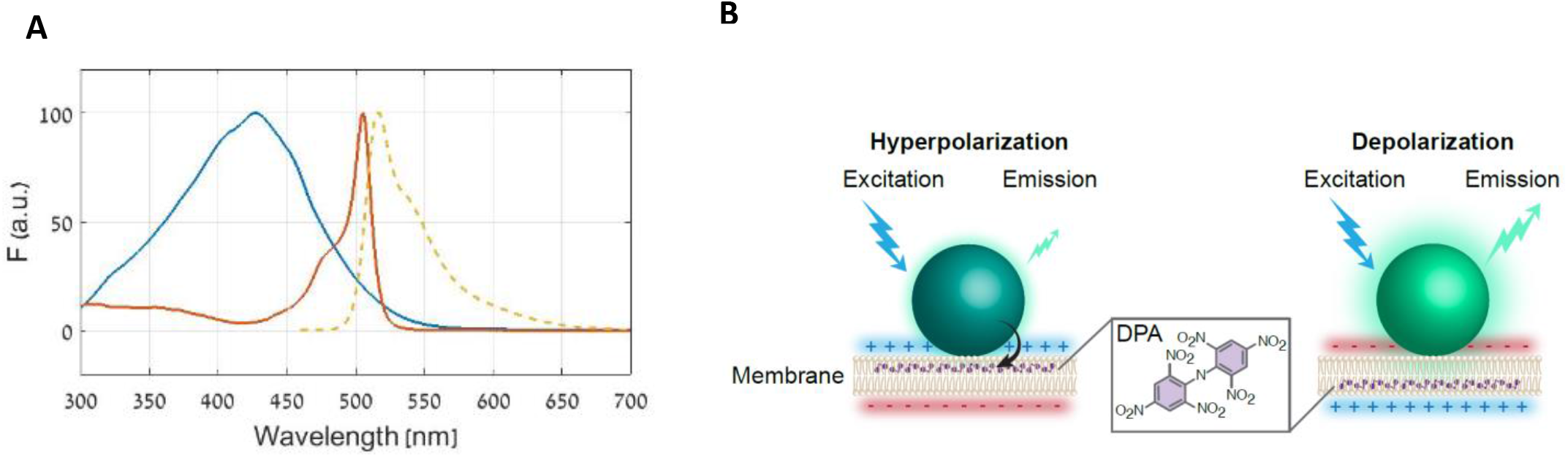
The FPS beads/DPA FRET pair mechanism. (A) Scheme of the fluorescent absorption and emission spectra; DPA absorption (blue), FPS Beads excitation (orange), FPS Beads emission (dashed yellow). An overlap of the FPS beads emission and DPA absorption affords FRET between the two species. (B) The FPS beads are attached to the extracellular side of the cell membrane; at hyperpolarization DPA molecules translocate to the outer leaflet of**_**the membrane and closer to the beads, resulting in FRET and a decrease in the bead’s fluorescence. Upon depolarization, the DPA molecules redistribute to the inner leaflet and away from the beads, resulting in an increase of the bead’s fluorescence.

The essence of the mechanism is as follows: The beads are applied to the bath solution and attach the cell membrane from the outside; the negatively charged DPA molecules in the membrane redistribute in response to the membrane potential. At negative potentials, the DPA molecules move closer to the outer leaflet of the membrane, resulting in quenching of the bead’s fluorophores. Conversely, upon depolarization, the DPA molecules move away from the beads toward the inner leaflet, so the fluorophores are unquenched (Fig.1B).

In this work, the potential of these FPS beads (with a hydrodynamic diameter of ~70 nm) as voltage nanosensors is explored. The performance of these beads as the donor particles to DPA is determined in HEK293 cells and primary cortical neurons. Fluorescence response to membrane potential changes is demonstrated from ensemble membrane staining as well as from single spots. In conclusion, this system’s sensitivity is adequate to perform membrane potential imaging from single particles, while from the aspect of kinetics, the temporal response is limited to ~5ms.

## Results

### FPS Beads and DPA comprise a membrane potential detector due to FRET reactions

As was detailed above, the FRET pair combined from DiO, a membrane dye, and DPA was described as a successful optical reporter for membrane potential [15], and was tested in our lab (Fig.SI1). In order to test the fluorescence response of the FPS beads/DPA FRET pair to membrane potential changes, simultaneous optical and electrical recordings were conducted. Isolated HEK-293 cells were labeled with beads, then the bath solution was washed and applied with DPA (see Materials and Methods). Using the whole-cell configuration of the patch-clamp technique, voltage steps were applied to the cell membrane. During the voltage protocol activation, the cells were excited with 485 nm and the fluorescence intensity was monitored (Fig. 2). When using the 60 mV steps voltage protocol described in Fig. 2C, upon depolarization the fluorescence intensity was increased with an averaged relative fractional change of 5.6 ± 2.4 % Δ*F/F* per 120 mV (n = 11) in the presence of 2 μM DPA in the bath solution. The labeling was stable after 2 hours in the bath solution, and patch-clamp experiments were still successfully applicable. Note that the fluorescence increases as the membrane voltage is more positive. This is in correlation with the beads/DPA FRET mechanism as described before. At positive membrane potentials the DPA molecules tend to be closer to the inner leaflet of the membrane, resulting in un-quenching of the beads fluorescence.

**Figure 2.**
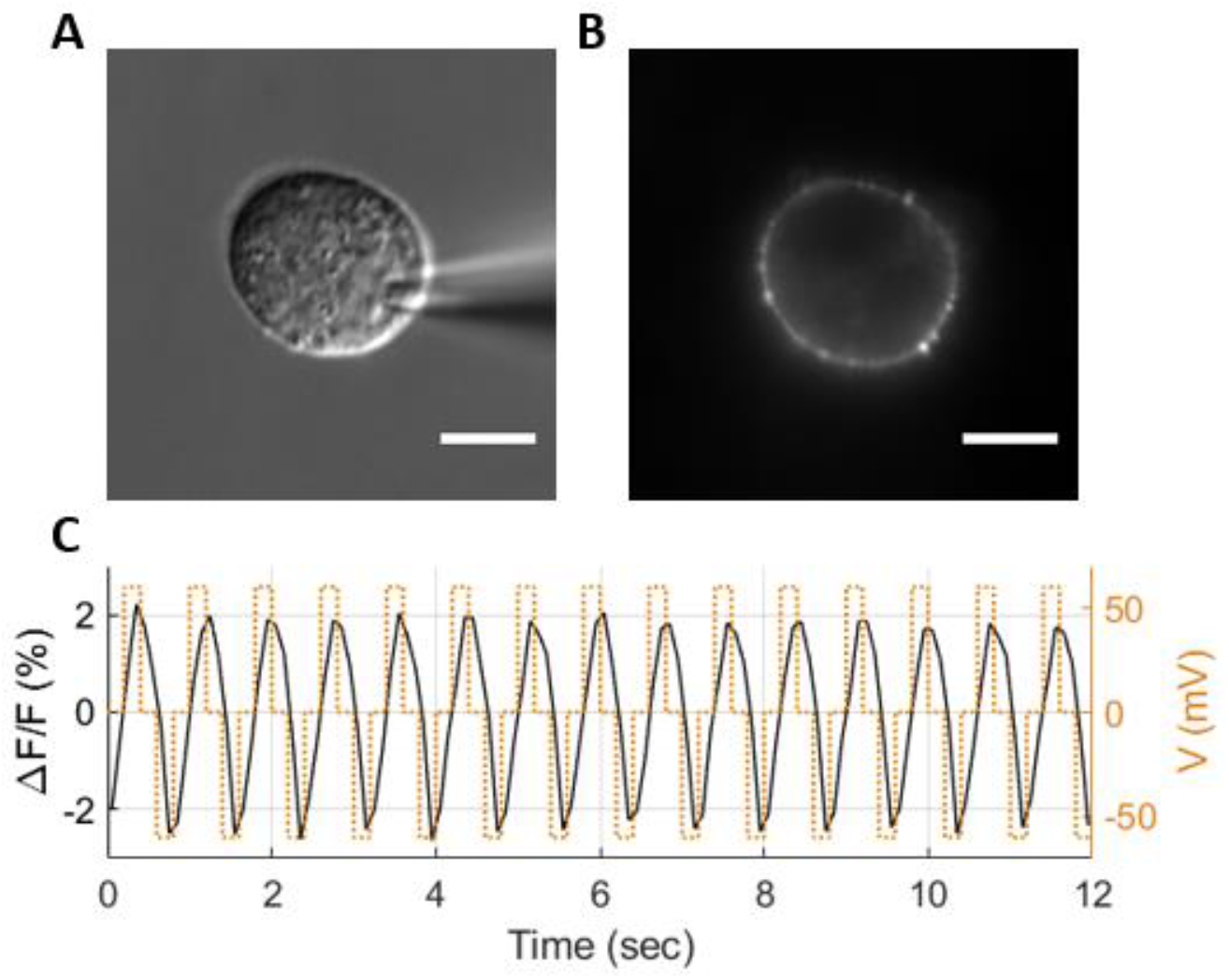
The fluorescence response of FPS beads/DPA FRET pair to membrane voltage in HEK cells. A DIC (A) and a fluorescence (B) image of a single HEK cell in the whole-cell patch-clamp mode, labeled with 0.5 nM beads, in the presence of 2 μM DPA in the bath solution. The patch electrode is seen in A. (C) The fluorescence response (black, solid) to membrane potential (orange, dashed) measured from the cell shown in A. The voltage was stepped from 0 mV to either +60 mV or −60 mV and back to 0 mV. The duration of each step was 200 msec, and 16 sec in total (12 sec are shown for better clarification). The average change in fluorescence emission ΔF/F from this cell was 4.3 ± 0.2 % per 120 mV. Scale bar: 10 μm.

The fluorescence response of FPS beads/DPA was further investigated. Different DPA concentrations (0.5 to 5 μM) were used, and the fluorescence response was tested for different membrane potentials (Fig. 3). As expected, the fluorescence intensity was higher in the presence of higher concentrations of DPA in the bath solution. At more positive membrane potentials tested, the %ΔF/F increased in higher DPA concentrations, while in lower concentrations it reached to a steady state. The averaged %Δ*F/F* at a membrane potential of +80 mV relative to the holding potential of −60 mV was 2.09 ± 0.34 % and 5.62 ± 1.28 % at 0.5 μM and 5 μM DPA, respectively (Fig. 3B). The averaged %Δ*F/F* at +60 mV and −60 mV relative to 0 mV (extracted from the 60 mV voltage-protocol as in Fig. 2C) in different DPA concentrations is also shown in Fig. SI2.Control experiments were also conducted in HEK cells. To confirm there is no significant response derived from FRET between DPA and endogenous proteins, voltage-clamp experiments were conducted in cells applied with 2 μM DPA, without the addition of beads (Fig. SI13A). The averaged Δ*F/F* per 120 mV was 0.14 ± 0.04% (n = 3), indicating there is no significant contribution to the beads/DPA response. Cells labeled with 0.5 nM beads only, without the addition of DPA (Fig. SI3B) were also tested. The averaged Δ*F/F* per 120 mV was 0.34 ± 0.14% (n = 12), which is a much smaller response than that of beads/DPA pair. A possible explanation for this response may be a change in ionic concentration at the vicinity of the bead’s fluorophores derived from the change in membrane potential.

**Figure 3.**
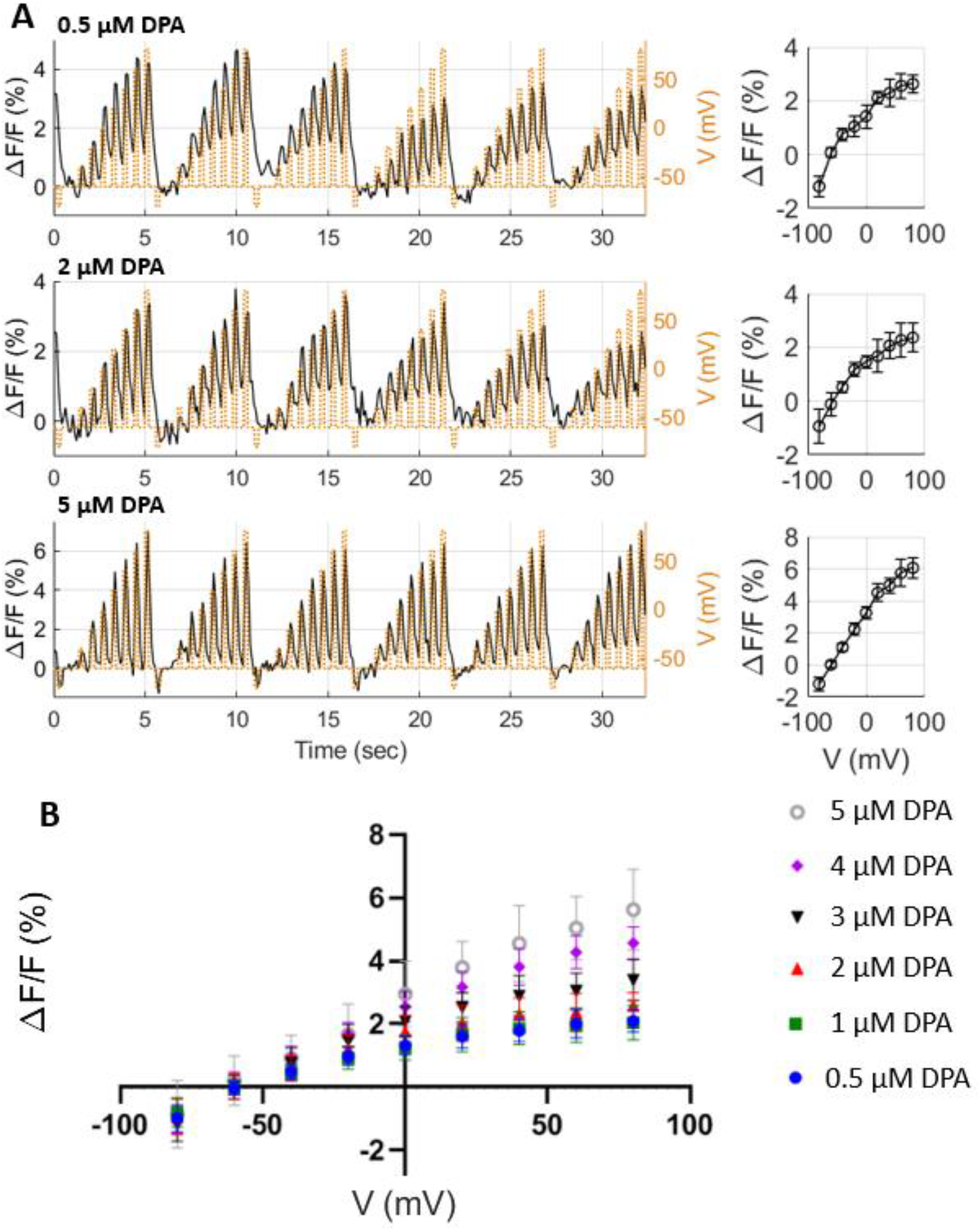
The fluorescence response of FPS beads/DPA in HEK cells for varying concentrations of DPA. The fluorescence responses of three cells in the presence of different concentrations of DPA are shown. The membrane potential was stepped in 20 mV increments from a holding potential of −60 mV. Each voltage level was held for 200 ms, followed by a repolarization step to −60 mV for 400 ms. Each trace was composed of six sweeps. The averaged fluorescence changes of these cells at each voltage relative to −60 mV are shown on the right. Error bars correspond to SD. (B) The averaged values of fluorescence changes to different membrane potentials in the presence of different concentrations of DPA (0.5 nM beads). The voltage protocol as in A. 5 μM DPA, n=6; 4 μM DPA, n=6; 3 μM DPA, n=7; 2 μM DPA, n=9; 1 μM DPA, n=5; 0.5 μM DPA, n=7. Error bars correspond to SD.

### Speed of FPS Beads/DPA fluorescence response to membrane potential changes

A previous publication reported a fluorescence time constant of ~ 0.1 ms in DiO/DPA FRET pair in HEK cells [15], similar to the speed of DPA movement in other preparations (~ 0.1 ms) [16, 17]. Another estimation of ~ 0.5 ms was reported in DPA and membrane tethered fluorescent proteins[18, 19].

To assess the temporal resolution of our system, the fluorescence response of FPS beads/DPA pair to voltage steps of −60 mV to +60 mV were measured in different pulse durations (5-100 msec) in HEK cells (Fig. 4). DPA concentration was determined to be 2 μM according to the experiments conducted with different DPA concentrations in HEK cells (Figs. 3 and SI2). A fluorescence response is clearly observed for voltage steps of 10 ms in duration (Fig.4B). At 5 ms pulses the fluorescence response is noticeable (Fig. 4), while no response is observed for shorter pulses. Thus, the temporal response of FPS beads/DPA is slower than was reported for other DPA FRET pairs (as described above), indicating this FRET pair is not suitable for measurements of AP-like signals. Since DPA translocates on the sub-millisecond time scale (This was also confirmed in our lab in experiments conducted with DiO/DPA in HEK cells; FigSI1), it is likely that the issue of slow response is related to the bead located on the membrane. In longer pulse durations, the system gets to a voltage-related steady-state as for the distance between the bead and DPA, while in short duration (<=5ms) it might not get to this state yet.

**Figure 4.**
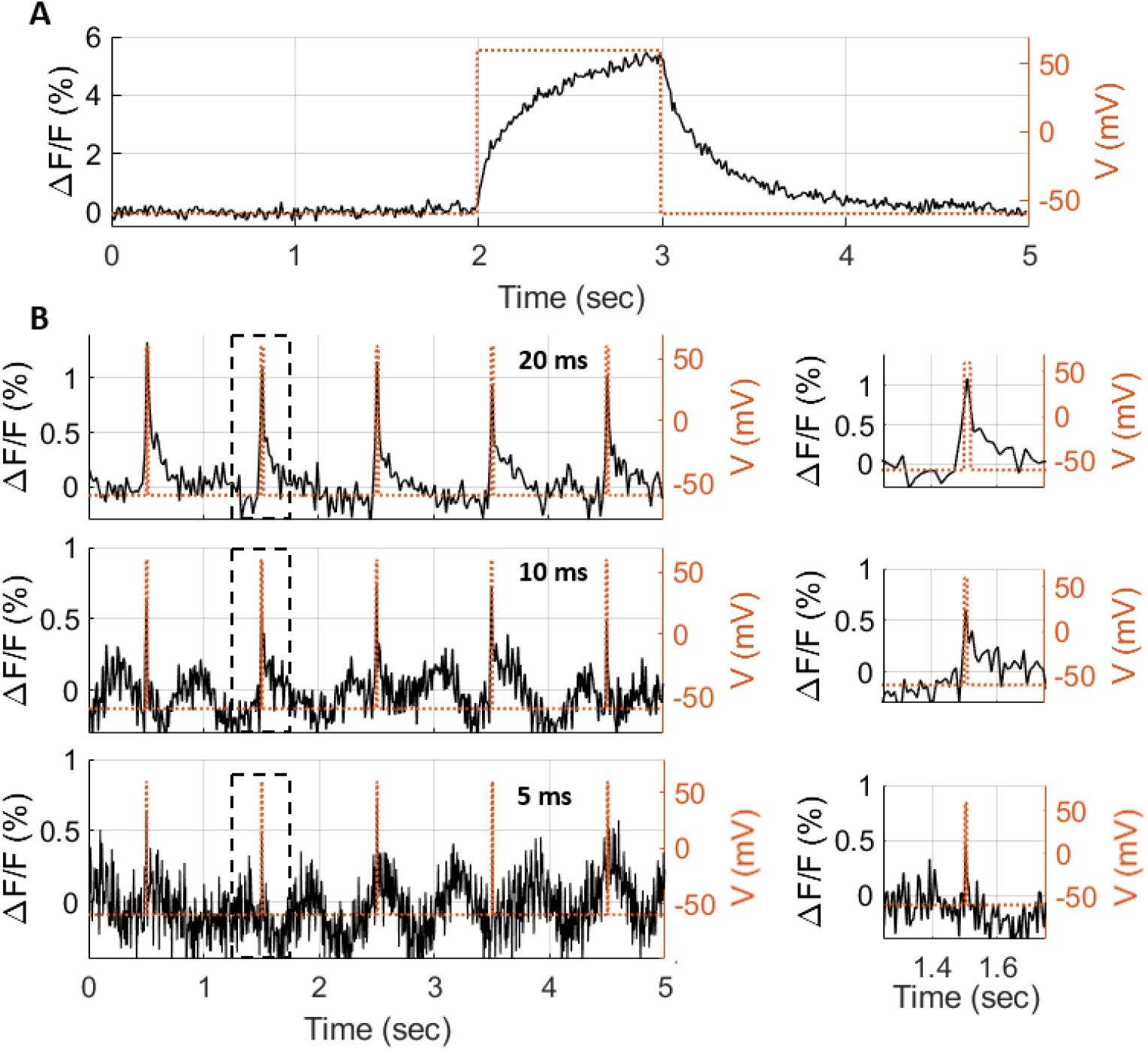
Response kinetics of FPS beads/DPA FRET pair to changes of membrane voltage in HEK cells. HEK cells were labeled with FPS beads and voltage-clamp experiments were conducted in the presence of 2 μM DPA. The membrane potential was stepped from a holding potential of −60 mV to +60 mV at different pulse durations, and repolarized back to −60 mV. (A) The change in fluorescence emission to 1 sec pulse, imaging at a frame rate of 100 Hz. A Slow response is observed at both the rising and falling parts of the polarization pulse. The response fall time, defined as the duration needed for the fluorescent emission to decay from 90% to 10% from its maximum intensity value once the pulsed polarization ends, was measured to be 670 ms. (B) The changes in fluorescence for 20, 10, and 5 ms pulses, imaging at a frame rate of 50, 100, and 200 Hz, respectively.

### Fluorescence response to membrane potential from single FPS Beads

A major advantage of using beads as voltage detectors over fluorescent dyes may be the potential of recording from single beads. Thus, our next goal was to test the FPS beads/DPA FRET mechanism in single beads imaging. Isolated HEK cells were labeled with 1 or 5 pM FPS beads to achieve a sparse staining (Fig. 5A). Cells that were labeled with one to six single fluorescent spots after DPA application were chosen for the experiment. The 60 mV steps voltage-clamp protocol as described in Fig. 2C was applied, and single ROIs were detected for analysis (Fig. 5A, right). Further description of spot detection analysis is described in materials and methods. Representative fluorescent traces of single spots are shown in Fig.5B. Only single spots that showed values larger than 2 for % Δ*F/F* per 120 mV were ascribed as responsive spots. In the presence of 2 μM DPA, 21 single spots out of 27 were responsive (77.8%), and showed an averaged Δ*F/F* of 3.48 ± 1.02% (measured from 8 cells). The presence of 5 μM DPA in the bath solution resulted in 12 responsive single spots out of 20 (60.0%), and showed a similar averaged Δ*F/F* of 3.33 ± 1.03% (measured from 5 cells) (Fig. 5C). Since the responsiveness of single spots in HEK cells was demonstrated, the fluorescence response of the FPS beads/DPA FRET pair to membrane potential was next examined in cultured neurons.

**Figure 5.**
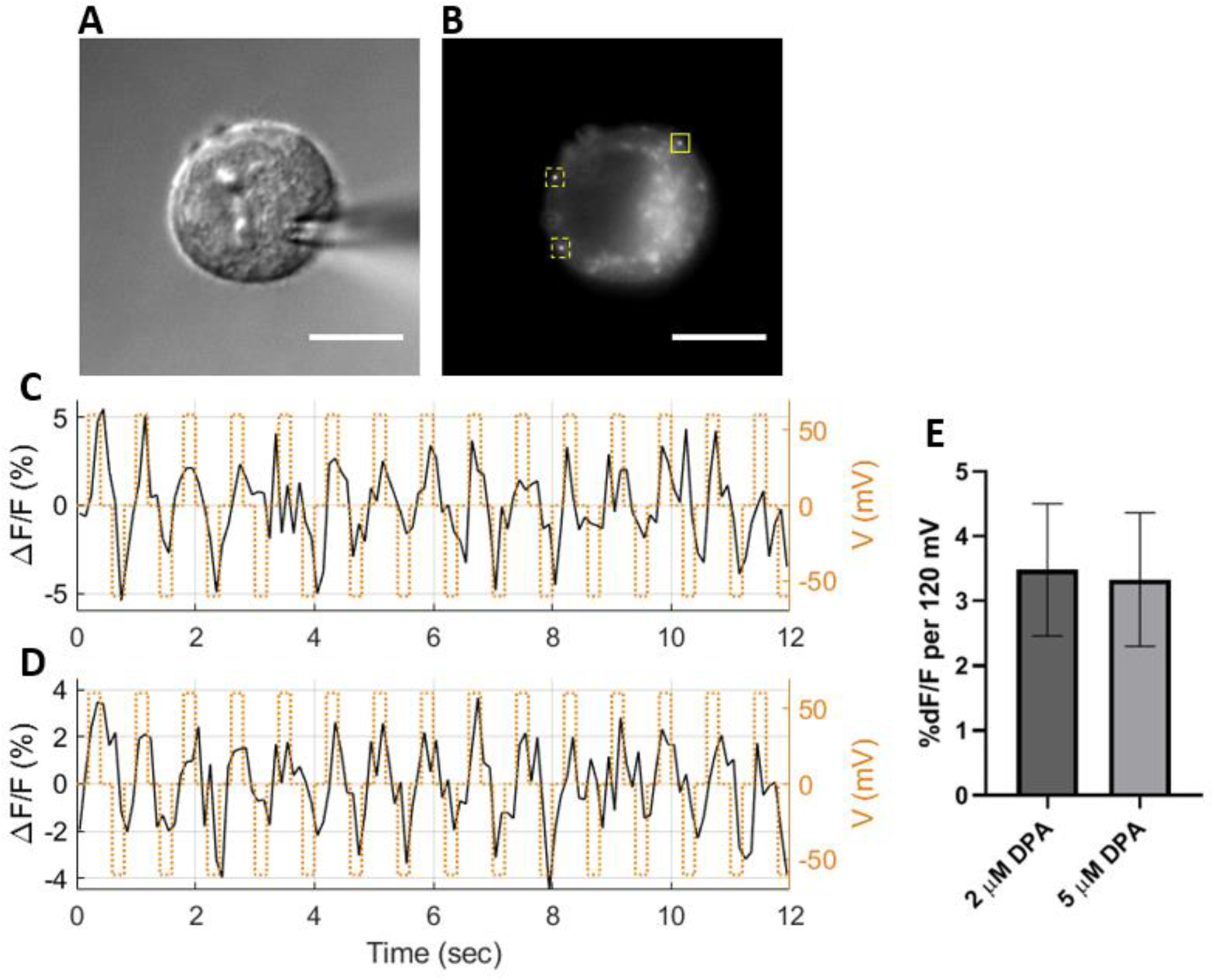
Optical responses to voltage of FPS beads/DPA FRET pair in HEK cells sparsely stained with beads. A DIC (A) and a fluorescence (B) image of a single HEK cell in the whole-cell patch-clamp mode, sparsely labeled with 5 pM beads, in the presence of 2 μM DPA in the bath solution. The voltage protocol is as described in fig. 2C. (C) and (D) show the fluorescence emission trajectory of one spot (indicated by the solid yellow square), and the averaged trajectory of three regions (indicated by yellow squares), respectively. The average change in fluorescence emission ΔF/F for the single spot and averaged trajectories was 4.5 ± 2.6 % and 4 ± 1.4 % per 120 mV, respectively. (E) The averaged change in fluorescence emission ΔF/F per 120 mV measured from single ROIs in cells exposed to 2 or 5 μM DPA in the bath solution. The cells were labeled with 1 or 5 pM beads. The voltage protocol is the same as in B. 2 μM DPA, n=21 measured from 8 cells; 5 μM DPA, n=12 measured from 5 cells. Error bars correspond to SD.

DPA is a lipophilic anion, as one it may effect excitable membranes by increasing their capacitance and change their electrical response properties [16, 19, 20]. To ensure the DPA concentration used in our experiments is suitable for cultured cortical neurons, whole-cell current clamp recordings were conducted in the presence of 1 μM DPA, 2 μM DPA or no DPA in the bath solution (Fig. 6). Action potentials (APs) were evoked by depolarizing current pulses in 50 pA increments (100-250 pA, 300 ms); No APs were evoked following 50 pA depolarizing currents. For consistency, the experiments of different DPA concentrations were conducted at the same DIV. These DPA concentrations had no significant effect on the first AP height, APs half width or threshold (P > 0.05); Fig.6 C-E).

**Figure 6.**
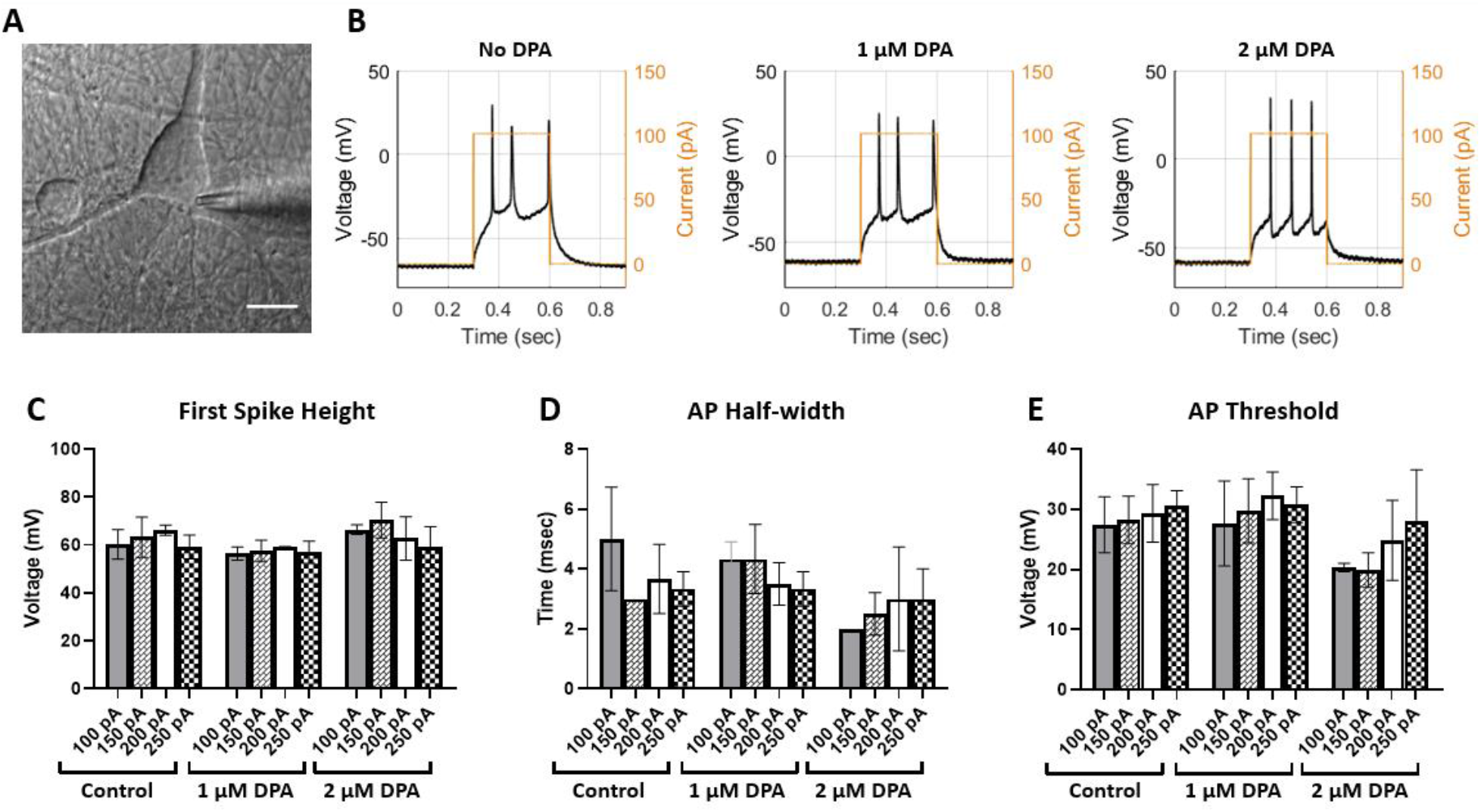
Electrical activity of cultured cortical neurons in the presence of different DPA concentrations. APs were recorded using the whole-cell configuration under the current-clamp mode, and were evoked by current injections. (A) A DIC image of a neuron in the whole-cell patch-clamp mode. The patch electrode is shown. (B) Spiking activity evoked by a square depolarization current pulse of 150 pA for 300 ms. The traces are attributed to neurons without DPA in the bath solution (left), in the presence of 1 μM DPA (middle) and 2 μM DPA (right). (C-E) Properties of the APs evoked by the protocol described in B. The APs properties are compared in different DPA concentrations and different amplitudes of depolarization current pulses. No significant difference was observed among columns and within columns (n= 3 for each treatment; P > 0.05; One-way-ANOVA).

Hence, the fluorescence response of FPS beads in neurons was tested in the presence of 2 μM DPA, the same concentration used in HEK cells and was already proven to be effective as a voltage detector in combination with beads.

Neurons were sparsely labeled with beads (22 pM), and single spots located on the neuron’s soma were detected and analyzed as described in materials and methods (Fig.7A). Representative fluorescent traces of single spots during application of 20 mV steps protocol are shown in Fig.7. The averaged %ΔF/F per 100 mV was 2.5 ± 1.0% for 18 single spots in 5 cells. These results demonstrate a correlation between the fluorescence of the bead’s single spots and membrane potential in cultures neurons. Thus, the principle of using FPS beads and DPA as a FRET-pair is successfully applicable for apparently single beads in neurons.

**Figure 7.**
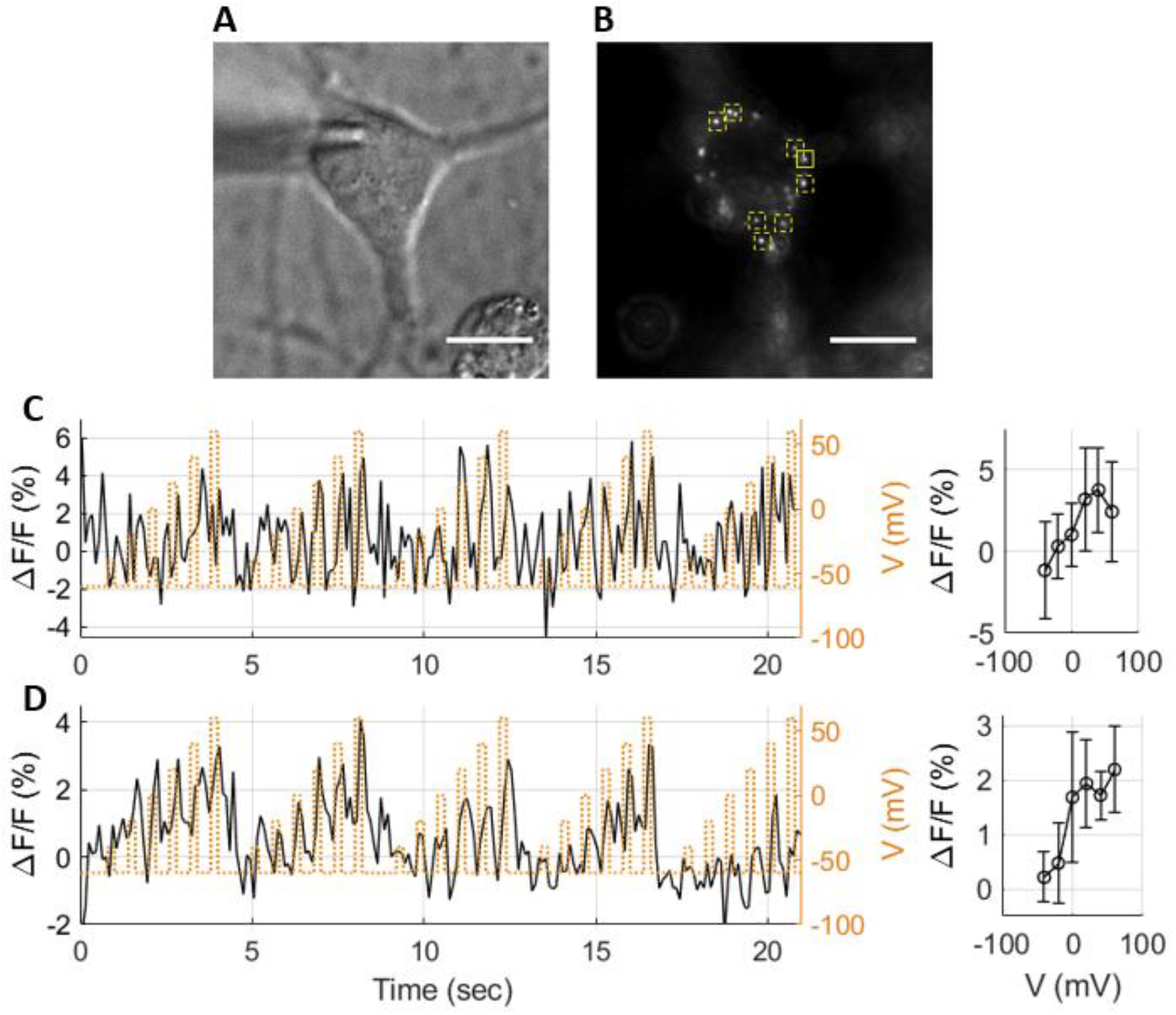
Optical responses to voltage of beads/DPA FRET pair in neurons sparsely stained with FPS beads. A DIC (A) and a fluorescence (B) image of a primary cortical neuron in the whole-cell patch-clamp mode, sparsely labeled with 22 pM beads, in the presence of 2 μM DPA in the bath solution. The membrane potential was stepped in 20 mV increments from a holding potential of −60 mV. Each voltage level was held for 200 ms, followed by a 400 ms repolarization step at −60 mV. Each trace was composed of five sweeps. (C) and (D) show the fluorescence emission trajectory (left) and average response (right) of one spot (indicated by the solid yellow square), and the average of nine spots (indicated by yellow squares), respectively. The average change in fluorescence emission ΔF/F for (C) and (D) was 3.2 ± 2 % and 2.2 ± 1.3 % per 100 mV, respectively. Error bars correspond to SD.

## Discussion

A new two-component system based on FRET between FPS beads, as the fixed fluorescent donors and DPA, a liphophilic anion which translocates between the leaflets of the membrane according to the membrane potential, is reported in this work. This system is demonstrated to correlate the membrane potential with fluorescent intensity in HEK cells and cultured neurons over a range of applied voltages. The correlation was confirmed not only for ensemble staining, but also, and more importantly, for single FPS beads.

This method has many required qualities over other approaches. First, the beads are applied to the extracellular solution, and adsorbed to the cell membranes spontaneously over a few minutes with no apparent cytotoxicity. According to the manufacturer, the beads surfaces are hydrophilic and negatively charged. No surface functionalization steps were needed for the adsorption of the beads to the cellular membranes. Note that the interaction of the beads with the membrane is not fully understood yet, and further characterization is needed. As for DPA, it has attractive properties as an acceptor, such as rapid kinetics(□<0.5 ms),hight voltage sensitivity within the physiological range, good aqueous solubility and low phototoxicity [15–17, 20]. DPA in the concentration used for recordings with FPS beads was also tested for potential effect on electrical properties of the neurons since it is located in the transmembrane electric field; No significant effect was shown for 2 μM DPA, as was also reported by Bradley et al. [15]. Secondly, the FPS beads demonstrate high photostability and brightness, which allows visualization and recordings from single particles. Altogether, in addition to the fact that the beads are commercial available in different sizes, and can be synthesized with functional surface groups, they constitute an easy approach for single particle voltage detection. The FPS beads can further be targeted to specific proteins via conjugation of recognition molecules, thus be targeted to specific cell types or sub-compartments. Future experiments are being planned in our lab to conjugate FPS beads with antibodies targeted to the extracellular domain of selected membrane proteins in cortical neurons. Considerations must be taken in choosing the right domain in order to minimize a potential interruption to the protein’s function.

Measurements of action-potentials demand a voltage-sensor displaying a fast submillisecond temporal response. When DPA is used with small organic donor dyes [21–24], the temporal response of the FRET-based system is adequate to capture action potentials [25]. However, the temporal response of FPS/DPA is estimated to be on the order of ~ 5ms (Fig. 4). This response is not suitable for capturing and recording action potentials. A similar slow temporal response was observed also with ND_TF,Cy5_ construct acting as membrane potential sensors on the nanoparticle level (DOI: 10.5281/zenodo.4704309). The observation of slow temporal response for two different types of nanoparticles points towards a similar underlying mechanism related to membrane adsorption. The reason for this slow temporal response of nanoparticles and for the slow response of the TF/DPA system is the subject of future studies Since DPA has rapid kinetics and DiO/DPA FRET pair, shows a fast sub millisecond response [15, 18, 19, 26, 27], it can be assumed that the speed limit is attributed to the FPS beads. Our hypothesis is that FPS/DPA temporal response could be improved by anchoring the beads to the membrane through cross-linking to several membrane proteins or by decorating the bead’s surface with long alkyl chains that can intercalate into the membrane. These approaches would be further investigated in order to maximize the FPS/DPA response in terms of kinetics.

In summary, the necessity of an optical tool for detecting voltage membrane changes in the single molecule level is undisputed. Such a tool that enables the investigation of local voltage membrane changes may contribute dramatically to the understanding of biological processes in neuroscience particularly. It can shed the light on the interactions between different parts of a neuron as well as the interactions between neurons in a neural network. Here, a new and accessible platform is demonstrated for membrane voltage imaging from single particles with a potential of targeting to specific desired membrane proteins.

## Supporting information

SI- Optical Probing of Local Membrane Potential with Fluorescent Polystyrene Beads

## Acknowledgments

This work has received funding from the European Research Council (ERC) under the European Union’s Horizon 2020 research and innovation program under grant agreement No. 669941, by the Human Frontier Science Program (HFSP) research grant RGP0061/2015, by the BER program of the Department of Energy Office of Science grant DE-FC03-02ER63421, and by the STROBE National Science Foundation Science & Technology Center, Grant No. DMR-1548924, by the Israel Science Foundation (ISF) Grant 813/19, by the Bar-Ilan Research & Development Co, the Israel Innovation Authority, Grant No. 63392.

## Data and code depository

Raw data fluorescent images of all voltage and electrical measurements for this paper can be found here: (https://zenodo.org, DOI: 10.5281/zenodo.4701597). Home-written code (MATLAB®) for extraction and analysis of recorded electrophysiological optical trajectories is available upon to request.

## Materials and Methods

### Cell line culture preparation

Human embryonic kidney (HEK293) cells were cultured in Dulbecco’s Modified Eagle Medium (DMEM) containing 10% fetal calf serum, 2 mM glutamine, 100 units/ml penicillin G and 100 μg/ml streptomycin, and grown in 5% CO2 at 37 °C under 90-95% humidity. For electrophysiology experiments the cells were seeded on glass coverslips (25-mm diameter) that were placed in 6-well plates and pre-coated with poly-D-lysine. Electrophysiology experiments were performed 24-48 hours after seeding with a cell confluence of 70%.

### Cortical neuron culture preparation

All animal experiments were approved by the local ethics committee for animal research (ethical approval number 63-09-2018). Primary cortical neurons were cultured from P1 and P2 newborn Sprague Dawley rats. The tissue was digested by 100 units of papain (Sigma-Aldrich) in Ca^2+/^Mg^2+^ free Hank’s balanced salt solution (HBSS) (Biological Industries, Israel) for 20 min in a 37° incubator. The tissue was then mechanically dissociated, and the cells were plated on 50 μg/ml poly-D-lysine-coated cover glasses in Neurobasal medium (Biological Industries, Israel) supplemented with 2% B-27 (Invitrogen), GlutaMAX-I 2mM (Invitrogen), Penicillin (100 unit/ml) / streptomycin (100μg/ml) (Biological Industries, Israel), 0.0055 Glucose and 5% Normal Horse Serum (Biological Industries, Israel). On the following day and twice a week thereafter, medium was exchanged with growth medium, containing the same components as described above, only the Neurobasal medium is exchanged with Minimum Essential Media (MEM). Once a week the medium was applied with 5μM Ara-C to prevent glial proliferation. Cells were incubated at 37 °C in a humidified 5% CO_2_ incubator. The experiments were conducted at 10-15 DIV.

### Staining cells with FPS beads and DPA

For HEK cells loading, 0.06 μm Carboxyl FPS yellow particles (Spherotech) were briefly sonicated and diluted right before use in Dulbecco’s Phosphate Buffered Saline (DPBS) to final concentrations of 500 pM for ensemble staining, and for 5 or 1 pM for single spots staining. DPA (20 mM in DMSO; Biotium) was diluted right before use in HEK external solution (detailed below). HEK cells seeded on glass coverslips were incubated with FPS beads for 10 minutes in room temperature (RT), washed, applied with DPA and immediately taken for imaging. For neurons loading, the particles were sonicated and diluted to final concentrations of 22 pM in neuronal external solution (detailed below). The cells were incubated for 5 minutes in RT, washed and applied with fresh DPA diluted in neuronal external solution.

### Electrophysiological recording

Voltage-clamp recordings in HEK cells were performed using the whole-cell configuration of the patch-clamp technique. The recordings were performed using a computer-controlled amplifier (EPC 10 USB, HEKA Elektronik). Recorded signals were filtered at 10 kHz using a Bessel filter, and digitized at 10 kHz to 100 kHz, depending on the timescale of the voltage waveform.

The imaging and electrophysiology experiments using HEK cells were conducted with the HEK external solution containing (in mM): 140 NaCl, 2.8 KCl, 2 CaCl_2_, 1 MgCl_2_, 10 HEPES and 10 glucose, adjusted with NaOH to pH 7.4 (310 mOsm). The pipette solution contained (in mM): 125 K-Gluconate, 0.6 MgCl_2_, 0.1 CaCl_2_, 1 EGTA, 10 HEPES, 4 Mg-ATP, 0.4 Na_2_GTP, adjusted with KOH to pH7.4 (295 mOsm). For experiments conducted in neurons, the neuronal external solution contained (in mM): 116.9 NaCl, 2.7 KCl, 1.8 CaCl_2_, 0.5 MgCl_2_, 20 HEPES, 5.5 glucose, 0.36 NaH_2_PO_4_, 200 Ascorbic Acid and Penicillin (100 unit / ml final) and Streptomycin (100 μg/ml final) at pH 7.3, and adjusted to 320 mOsm with sucrose. The neuronal pipette solution contained (in mM): 110 K-Gluconate, 10 KCl, 10 Glucose, 8 Na-Creatine-phospate, 5 EGTA, 10 HEPES, 4 Mg-ATP, 0.4 Na-GTP (pH 7.3); adjusted to 290 mOsm with sucrose. The patch pipettes were pulled from borosilicate glass (Warner Instrument) with a resistance of 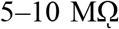 when filled with the pipette solution.

### Optical Recordings

Optical recording. Imaging was performed using an LED excitation (SPECTRA X, Lumencor) passing through a 470/24 nm bandpass filter, and an EMCCD camera (iXon Ultra 897, Andor), coupled to a motorized inverted microscope (IX83, Olympus). All imaging was performed using a 100X oil-immersion objective (UAPON100XOTIRF), apart from the data shown in Fig. 3, where a 60X oil-immersion objective (UPLSAPO60XO) was used. Fluorescence emission was filtered with a bandpass filter (ET520/20m, Chroma). A DAQ device (USB-6211, National Instruments), controlled by a home-written software (LabVIEW, National Instruments), was used for synchronizing the camera and the patch clamp amplifier.

### Data Analysis

Analysis of fluorescence imaging of clamped cells was performed in MATLAB (MathWorks). For sparse staining, a 2D particle tracking algorithm was used in order to correct for small translations. Fluorescence intensity for each frame was calculated by averaging the 10 brightest pixels of each spot for sparse staining, and averaging the fluorescence from a whole cell for ensemble measurements. For the latter, background-subtraction of each frame was performed by subtracting the mean value of the 10% darkest pixels. Fluorescence trajectories were corrected for photobleaching by either a shortpass filter or subtraction of the fluoresence trajectory by a two-term exponential function fitted to the baseline voltage levels. The average optical voltage response for each intensity trace was calculated by averaging the intensity value corresponding to every voltage level. In order to achieve a stabilized value of the response, the averaging was performed on the frame containing the last 100 ms of each voltage step.

