## Supplementary material for "Optical Probing of Local Membrane Potential with Fluorescent Polystyrene Beads": SI- Optical Probing of Local Membrane Potential with Fluorescent Polystyrene Beads

### Supplemental Information

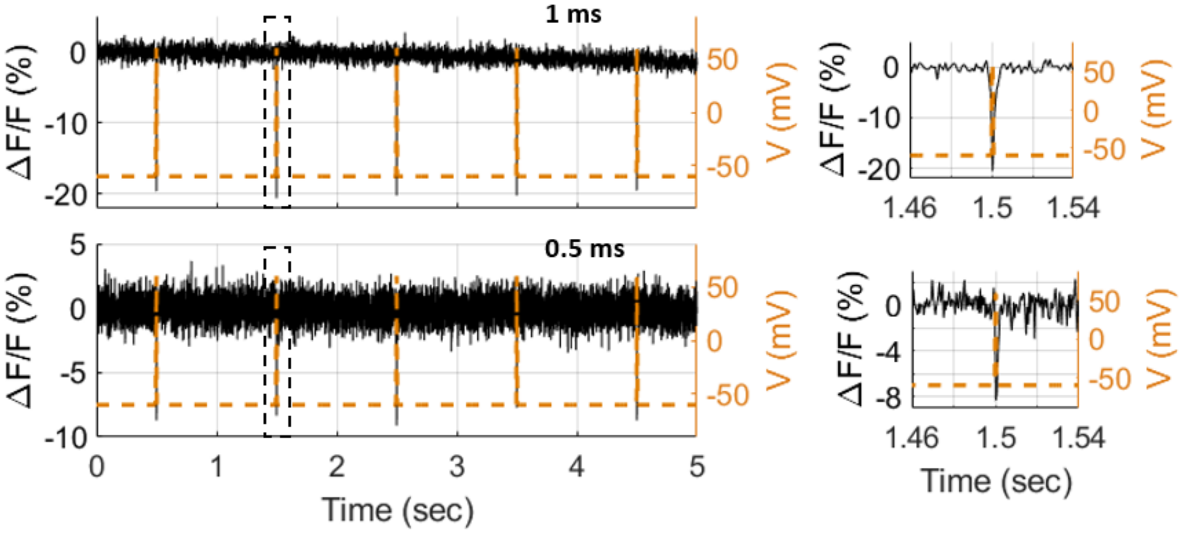


**Figure SI1. Response kinetics of DiO/DPA FRET pair to changes of membrane voltage in HEK cells.** HEK cells were labeled with 10 µM DiO, and the wh and voltage-clamped in the presence of 2 µM DPA to a holding potential of -60 mV, with transient polarization pulses to +60 mV at different durations. **(B)** The changes in fluorescence for 1 and 0.5 ms pulses, imaging at a frame rate of 1 and 2 kHz, respectively.

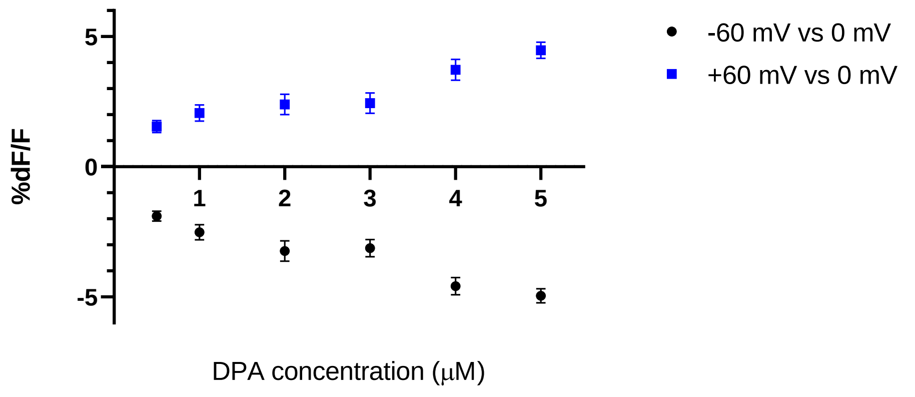


**Figure SI2. The averaged fluorescence response of FPS beads/DPA to membrane voltage in the presence of different concentrations of DPA.** HEK cells were labeled with 0.5 nM beads, and the bath solution was applied with 0.5 to 5 µM DPA. Whole-cell patch clamp was carried out and the membrane voltage was stepped as described in Fig.2C. The averaged %Δ*F*/*F* at +60 mV and -60 mV are shown. Error bars correspond to SD. 5 µM DPA, n=6; 4 µM DPA, n=6; 3 µM DPA, n=6; 2 µM DPA, n=11; 1 µM DPA, n=5; 0.5 µM DPA, n=7.

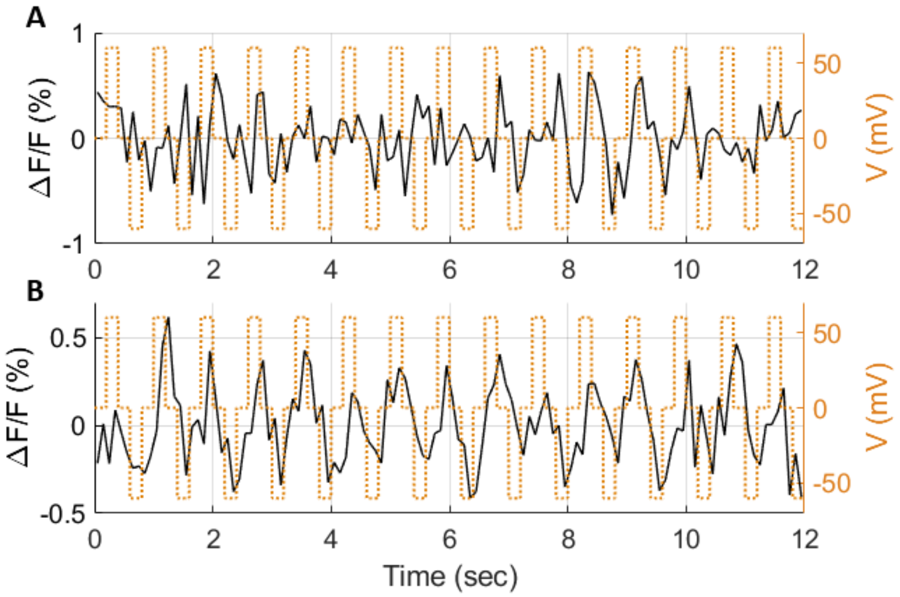


**Figure SI3**. **Control fluorescence response measurements in HEK cells.** Representative optical response (black, solid) to changes in membrane potential (orange, dashed), for cells which were **(A)** exposed to 2 µM DPA only, without the addition of beads, and (**B)** labeled with 0.5 nM beads without the addition of DPA. The voltage protocol is as described in fig. 2C. The average change in fluorescence emission ΔF/F for DPA-only and bead-only labeling was 0.2 ± 0.5 % and 0.5 ± 0.2 % per 120 mV, respectively.
